# N-acetyl-L-leucine (Levacetylleucine) normalizes Transcription Factor EB (TFEB) activity by stereospecific bidirectional modulation

**DOI:** 10.64898/2025.11.30.691375

**Authors:** Lianne C. Davis, Rebecca Braine, Grant C. Churchill, Mallory Factor, Taylor Fields, Marc Patterson, Frances Platt, Michael Strupp, Antony Galione

## Abstract

Levacetylleucine (Aqneursa^TM^), a chemically modified amino acid, is the only US Food and Drug Administration-approved monotherapy for the treatment of Niemann-Pick disease type C (NPC) (Beninger, 2024; Mullard, 2024; van Gool et al., 2025). This acetylated derivative of L-leucine functions as a pro-drug, with the acetyl group rendering it a substrate for the monocarboxylate transporter (MCT) family of transporters to allow appreciable penetration of the blood-brain barrier and its efficient uptake into cells (Churchill et al., 2021). Inside cells, levacetylleucine undergoes metabolism catalysed by acylases, and the resultant high quantities of L-leucine enter metabolic pathways which enhance mitochondrial bioenergetics and, as previously demonstrated, indirectly ameliorate lysosomal function (Kaya et al., 2020). Here, we show a novel aspect of levacetylleucine’s mechanism of action, demonstrating a direct effect on lysosomal function through its rapid modulation of the translocation of the transcription factor TFEB, a master regulator of lysosomal biogenic and autophagic genes (Napolitano and Ballabio, 2016), from cytoplasm to nucleus. Uniquely, we have demonstrated a biphasic action whereby levacetylleucine normalizes TFEB activity, consistent with levacetylleucine’s previously shown ability to regulate cellular homeostasis: in wild-type HeLa cells, levacetylleucine enhances and activates the translocation of TFEB to the nucleus. In contrast, in cellular models of NPC type 1 disease, where TFEB is already over-expressed in the nucleus (as the cell attempts to compensate for the primary defect by activating TFEB as a natural cellular response to the lysosomal substrate accumulation and associated cellular stress), treatment with levacetylleucine down-regulates and restores the distribution of TFEB to a more normalized cytoplasmic: nuclear ratio. Importantly, both effects of levacetylleucine occur at concentrations consistent with plasma concentrations in therapeutic dosing (Churchill et al., 2020). The effects were also confirmed to be stereospecific to the L-enantiomer, as neither the D-enantiomer (N-acetyl-D-leucine) or racemate (N-acetyl-DL-leucine) had any effect, The presence of the D-enantiomer in the racemic mixture inhibited the ability of levacetylleucine to promote TFEB bidirectional translocation, consistent with previous studies, which have established antagonism of N-acetyl-L-leucine by N-acetyl-D-leucine in the racemic mixture (rendering the racemic mixture without effect). This bidirectional mechanism of action of levacetylleucine to impact lysosomal function directly and normalize, either by activating basal TFEB signalling or reducing aberrant TFEB function in NPC1 knockout cells, thereby modulating lysosomal and autophagic functions, lends itself to the treatment of a broad range of neurological and neurodevelopment disorders.

## Introduction

Levacetylleucine (Aqneursa^TM^), an orally-administered amino acid derivative, is the only US Food and Drug Administration (FDA) approved monotherapy for the rare genetic lysosomal storage disease (LSD), Niemann-Pick disease type C (NPC), conferring rapid symptomatic relief (Bremova-Ertl et al., 2024) as well as long-term disease-modifying effects (Patterson et al., 2025). The parent molecule, L-leucine, is a zwitterion at physiological pH, and is transported into cells by the easily saturable transporter L-type amino-acid transporter (LAT1). Addition of the acetyl moiety to L-leucine confers a net negative charge to the molecule at physiological pH, allowing levacetylleucine to be taken up into cells by high-capacity monocarboxylate transporters (MCTs), which are ubiquitously expressed, thereby delivering the drug to all tissues, including the central nervous system, by readily crossing the blood-brain barrier (Churchill et al., 2021). Inside cells, levacetylleucine enters enzyme-controlled pathways that correct metabolic dysfunction and enhance energy (ATP) production, which also leads to an improvement in lysosomal function (Kaya et al., 2020; Kaya et al., 2021). This has multiple consequential effects: mitochondrial and lysosomal function are intrinsically linked, and interact at membrane contact sites (Martello et al., 2020) and the normalization of energy metabolism improves lysosomal function, leading to a reduction in the storage of unesterified cholesterol and sphingolipids (Kaya et al., 2021). In addition, levacetylleucine corrects aberrant membrane contact sites in NPC patient cells where there is a deficit of endoplasmic reticulum (ER)-lysosomal contacts for efficient lipid transfer (Hoglinger et al., 2019) and a concomitant inappropriate gain of mitochondrial-ER contacts with resultant mitochondrial lipid accumulation (Kiraly, 2025).

In various animal models, levacetylleucine treatment shows a slowing of neurodegeneration and also leads to a dampening of neuroinflammation, consistent with the drug’s neuroprotective effects (Hegdekar et al., 2021; Kaya et al., 2021): the latter was also shown in patients with NPC (Dinkel et al., 2024). Due to its multi-modal mechanism of action, levacetylleucine has potential and is being developed for a range of rare and common neurodegenerative and neurodevelopmental disorders.

Niemann-Pick disease type C is a rare, serious, debilitating, pre-maturely fatal disorder affecting 1/100,000 live births (Tifft, 2024), and one disease in a family of over 70 rare monogenic LSDs that collectively affect 1:5000 live births. NPC is caused by mutations in either the *NPC1* or *NPC2* genes. The proteins they encode, NPC1 or NPC2, respectively, function cooperatively to facilitate lipid transport from the lysosome to the ER (Hoglinger et al., 2019). The NPC1 protein is a transmembrane protein involved in lipid transport, whilst NPC2 is a soluble luminal lysosomal protein thought to be involved in transferring lipids to NPC1 (Rosenbaum and Maxfield, 2011). NPC gene dysfunction leads to the accumulation of multiple classes of lipids in lysosomes, causing lysosomal enlargement, disruption of intracellular lipid trafficking and autophagy, dysregulation of mTOR signalling, reduction in lysosomal Ca^2+^ storage and release, perturbation of energy metabolism and mitochondria function, and ultimately cell death (Platt et al., 2018). Although all organs are affected, neurons are particularly susceptible with symptoms manifesting as impaired motor function and cognition.

Recently, it has been demonstrated that small molecule activation of transcription factor EB (TFEB) is impaired in NPC1^-/-^ cells, contributing to impaired lysosomal clearance (Du et al., 2025). TFEB, a key transcription factor affecting lysosomes, orchestrates the expression of genes (the coordinated lysosomal expression and regulation (CLEAR) network) involved in lysosomal biogenesis and autophagy, as it promotes the expression of genes required for autophagosome formation, lysosome biogenesis, and lysosomal function and exocytosis (Napolitano and Ballabio, 2016). TFEB contains basic helix-loop-helix-leucine zipper domains (bHLH-Zip) and belongs to the microphthalmia MiT-TFE family of transcription factors (Puertollano et al., 2018), and is highly expressed in the CNS (Cortes and La Spada, 2019). In its inactive phosphorylated state, TFEB resides in the cytoplasm; upon dephosphorylation it dissociates from 14-3-3 proteins and translocates to the nucleus to regulate gene expression (Rosenbaum and Maxfield, 2011). Dysregulation of TFEB activity is implicated in various neurodegenerative and neurodevelopmental diseases (Chen et al., 2024). For instance, previous studies have demonstrated that targeting the TFEB pathway has neuroprotective effects in various *in vivo* or *in vitro* models of Alzheimer’s disease. Hence, small-molecule TFEB activators or exogenously overexpressed TFEB may promote lysosomal function and autophagic flux and may prevent, obstruct, or reverse the pathogenesis of neurodegenerative diseases (Gu et al., 2022).

In view of its effects on a broad range of NPC disease manifestations referable to compromise of the lysosomal-mitochondrial axis, we investigated the role of levacetylleucine in modulating TFEB activity, the key regulator of this final common pathway of neurologic dysfunction.

## Methods

### Cell Culture and Transfection

HeLa cells were cultured in DMEM supplemented with 10% v/v FCS, 2 mM glutamine, 100 U/ml penicillin and 100 µg/ml streptomycin, at 37°C under 5% CO_2_. Cells were trypsinized and seeded onto CellView Slides (Greiner Bio-One).1-2 days after sub-culturing, cells were transiently transfected with JetPEI reagent in a 5:2 ratio with DNA. Per well, cells were transfected with 100 ng TFEB tagged on its C-terminus with either EGFP (Addgene plasmid # 38119) or mScarlet3 (produced in-house) for 4-6 hrs. Transfection medium was removed and replaced with fresh DMEM with or without agents and incubated overnight at 37°C. The next day, TFEB-EGFP-expressing cells were loaded for 1h at 37°C with NucSpot® Live 650 Nuclear Stain (Biotium) in the continued presence of drugs as required. Cells were then transferred into extracellular medium (ECM, mM: 121 NaCl, 5.4 KCl, 0.8 MgCl2, 1.8 CaCl2, 6 NaHCO3, 25 HEPES, 10 Glucose) that maintained the overnight treatment reagents and these live cells were imaged immediately.

Cells expressing TFEB-mScarlet3 were treated with similar reagent protocols but could be fixed with 4% PFA (in PBS) after reagent treatments. After fixation and permeabilization (0.1% Triton X-100 in PBS for 15 mins), nuclei were labelled with NucSpot® Live 488 Nuclear Stain (Biotium) for 10 mins in PBS and imaged within 2 days.

### Microscopy

Cells were imaged at room temperature using a Nikon A1R laser-scanning confocal equipped with a Plan ApoVC 20x DIC N2 (NA: 0.75) or Plan Fluor 40x oil DIC H N2 (NA: 1.3) objective. In Channel-Series mode, green, red or far-red fluorophores were alternately excited (ex/em): 488/525 nm, 561/595 nm, and 640/700 nm respectively.

### Nuclear Translocation Analysis

To quantify the translocation of fluorescent TFEB from the cytoplasm to the nucleus, we analysed the Pearson’s Correlation Coefficient between TFEB and the orthogonal nuclear stain in single cells. Cells with a cytoplasmic location have negative coefficients, whereas nuclear translocation is reflected by positive coefficients. Cells with nuclear TFEB (partial or completely translocated) were defined as having a Pearson’s coefficient >0, and the number of cells satisfying this criterion expressed as a percentage of the total number.

### Reverse Transcription Quantitative PCR (RT-qPCR)

Total RNA was extracted from wild-type HeLa cells using a RNeasy Plus Mini kit (Qiagen) following the manufacturer’s instructions: quality and quantity were assessed with a NanoDrop spectrophotometer (Thermo Fisher Scientific). cDNA was prepared using an iScript cDNA synthesis kit in a MyCycler Thermal Cycler system (both Bio-Rad). qPCR was then performed with a CFX96 Real-Time PCR instrument (Bio-Rad) using PowerUp SYBR Green (Applied Biosystems) and the primer pairs specified in **Table 1**. (sourced from OriGene). Threshold cycle (C_t_) values were normalised to the reference gene ß-actin using the comparative C_t_ method, on a scale where β-actin expression equals 10,000 units.

**Table 1.**
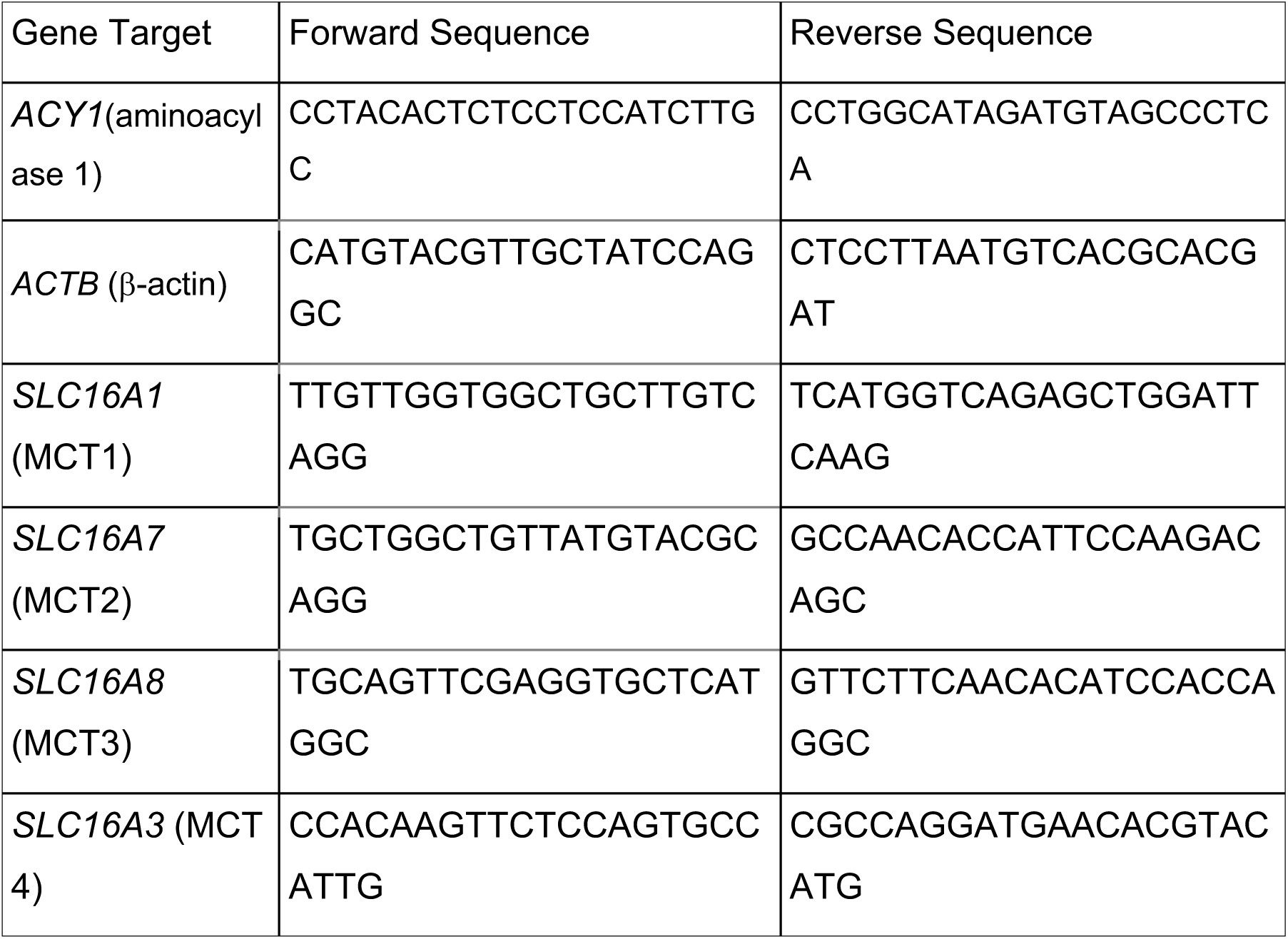
Primer sequences for RT-qPCR, sourced from OriGene and PrimerBank.

## Results

### Expression pattern of monocarboxylate transporters and aminoacylase 1 enzyme

We have previously proposed that levacetylleucine acts as a pro-drug to generate high concentrations of L-leucine in cells since it is a substrate for MCT transporters (Churchill et al., 2021). Therefore, in the first series of experiments we investigated whether HeLa cells express the key components of the levacetylleucine transport mechanisms previously proposed. We found that HeLa cells express the transporter system and acylase that are thought necessary for a response to levacetylleucine. Expression patterns of monocarboxylate transporters (MCT1, MCT2, MCT3, MCT4) and the aminoacylase 1 enzyme (ACY1) were determined by RT-qPCR (primer sequences in **Table 1**). Expression was normalized to β -actin, on a scale where β -actin expression equals 10,000 arbitrary units. We show that HeLa cells contain transcribed mRNA for three monocarboxylate transporter types, including MCT1 1, (**Fig. S1**) as well as an acylase shown to hydrolyze N-acetyl groups from amino acids (Bloch and Borek, 1946; Bloch and Rittenberg, 1947). The resulting L-leucine enters metabolic pathways to increase ATP synthesis, but also acts to clear stored lipids from lysosomes (Kaya et al., 2020).

### Effects of levacetylleucine, acetyl-D-Leucine, the racemate and L-leucine on translocation of TFEP-GFP and its functional consequences

To test for a direct effect of the drug on lysosomal function, we examined the effect of treating HeLa with levacetylleucine on the translocation of TFEB-GFP from cytoplasm to nucleus, a hallmark for the activation of this transcription factor (Settembre et al., 2012). We found that extracellular incubation of cells with levacetylleucine (2 mM) caused the appearance of TFEB-GFP in the nucleus during an 18-hour incubation (**Fig. 1A**). The effect was concentration-dependent with effects seen at the sub-millimolar range (**Fig. 1B, C**), which coincides with plasma concentrations observed in mice dosed with therapeutic concentrations of levacetylleucine (Churchill et al., 2020).

**Figure 1.**
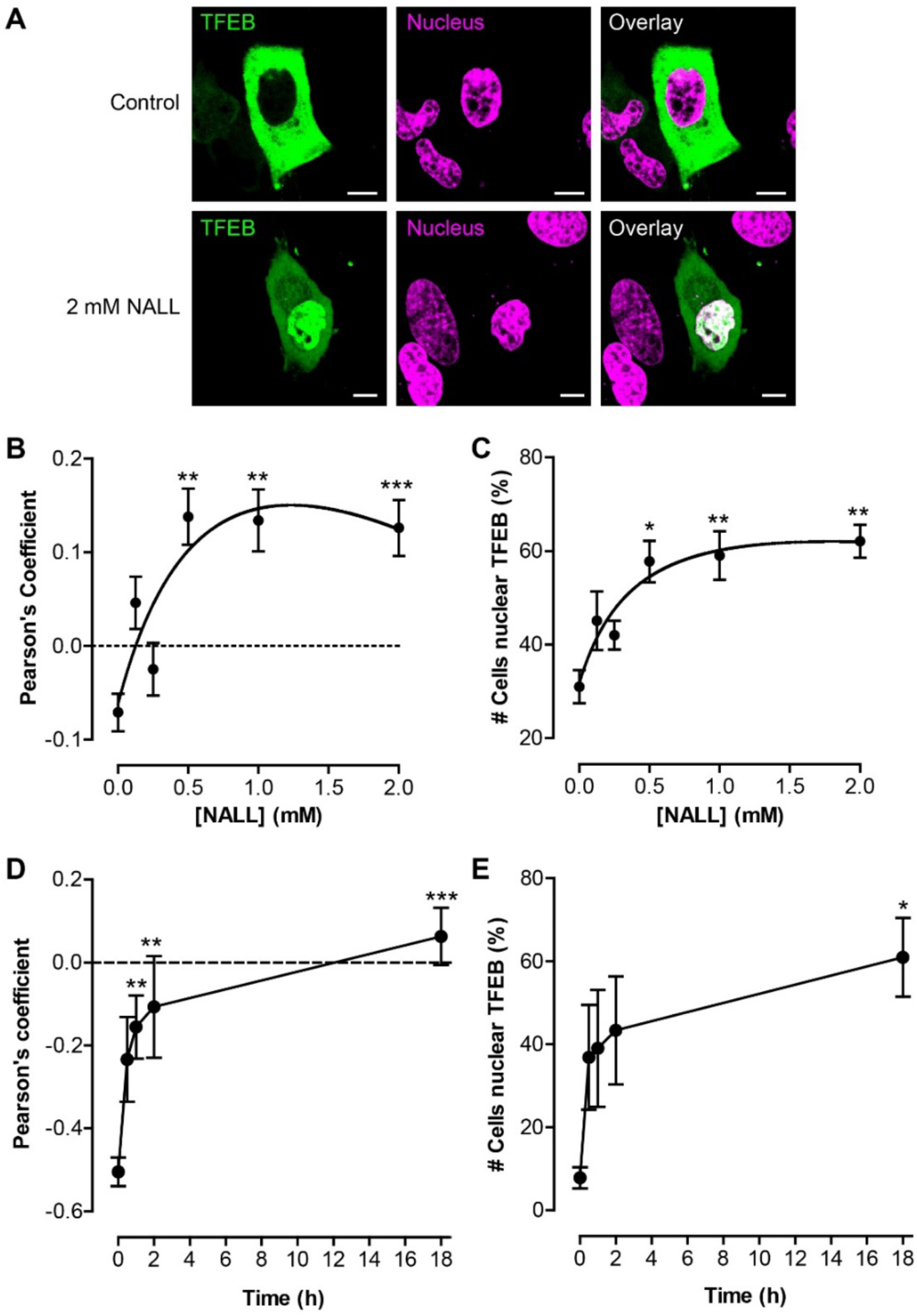
NALL induces nuclear translocation of TFEB. HeLa cells were transfected with TFEB-EGFP (A-C) or TFEB-mScarlet3 (D,E) and then incubated with NALL at different concentrations or times. Cells were then counter-stained with the NucSpot 488 (D,E) or NucSpot 650 (A-C) to label the nucleus and TFEB translocation quantified by colocalization with NucSpot. (A) Single-cell images of TFEB-EGFP (green) and NucSpot 650 (magenta). (B,C) Concentration-response of HeLa treated for 18 hrs with NALL. Nuclear localization is plotted as either the Pearson’s coefficient (B) or derived as the percentage of cells with nuclear TFEB (C). (D,E) Time course of the effect of 2 mM NALL upon TFEB, either expressed as the Pearson’s coefficient (D) or the percentage of cells with nuclear TFEB (E). The dotted lines (B,D) highlight a correlation coefficient of zero used for thresholding the percentage of nuclear cells. Data are expressed as the mean ± SEM of 179-215 cells, and significance determined by ANOVA test, with significance depicted as * *p* < 0.05, ** *p* < 0.01, or *** *p* < 0.001 compared to respective control, 0 mM (B,C) or t = 0 (D,E).

The onset of action of levacetylleucine was rapid, with half-maximal effect seen within 60 min (**Fig. 1 D, E**) during continuous application of the drug. Importantly, this effect was not shared by L-leucine over a similar concentration range (**Fig. 2**), which is the major metabolite of levacetylleucine through the action of amino acid acylases (Birnbaum et al., 1952), demonstrating that this effect required the acetyl form of leucine (as in levacetylleucine). This effect is likely only be transient since levacetylleucine is converted to L-leucine by cellular acylases (Bloch and Rittenberg, 1947; Churchill et al., 2020).

**Fig. 2.**
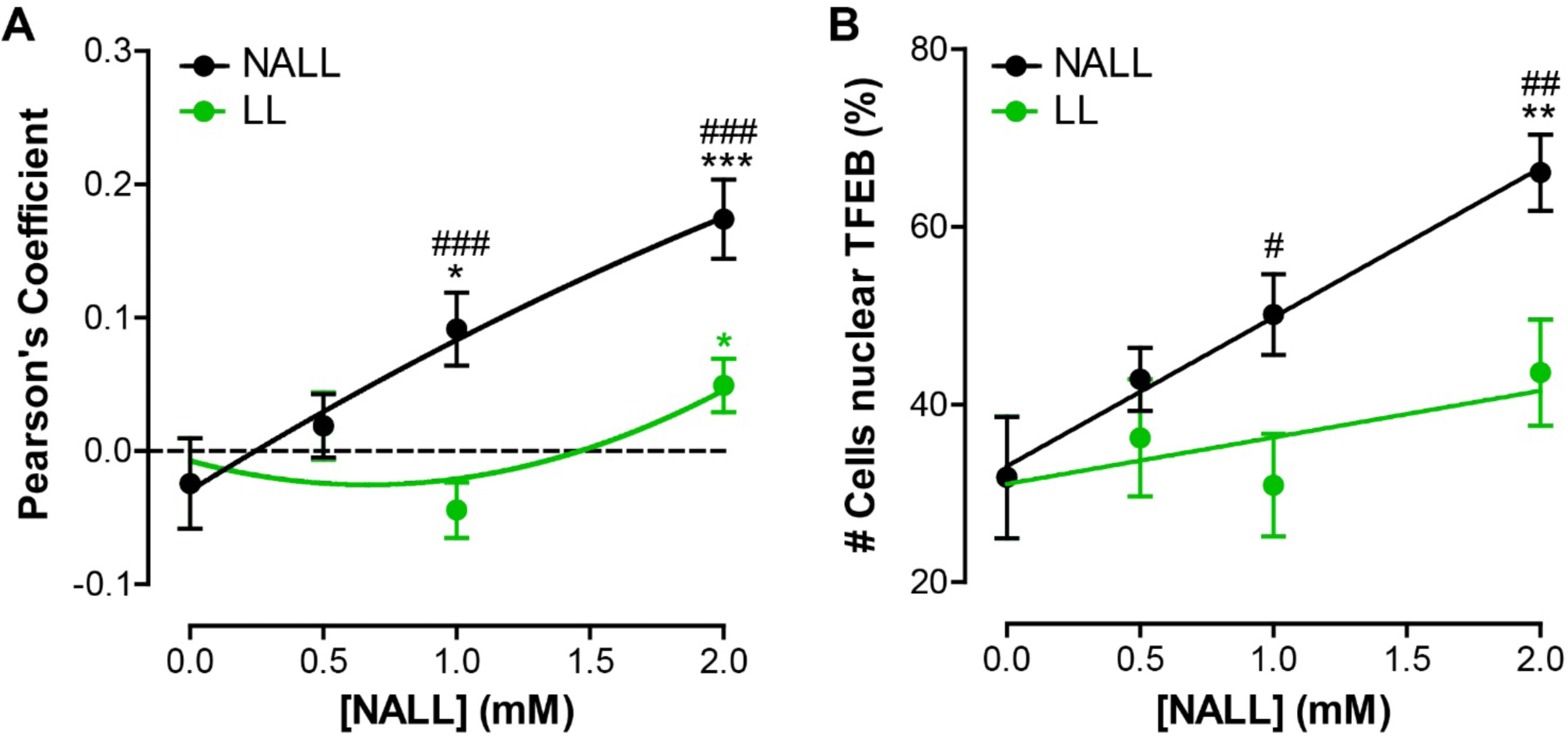
NALL induces TFEB translocation more efficaciously than does L-leucine. TFEB-EGFP translocation in HeLa cells incubated for 18 hrs with different concentrations of either NALL (black) or L-Leucine (LL, green). Translocation is expressed either as the Pearson’s correlation coefficient with NucSpot 650 (A) or as the percentage of cells with nuclear TFEB (B). Data are expressed as the mean ± SEM of 104-161 cells, and significance determined by nonparametric ANOVA test (A) or unpaired t-test (B), with significance depicted as * *p* < 0.05, ** *p* < 0.01, or *** *p*< 0.001 compared to the vehicle control, i.e. 0 mM. Comparing NALL and LL: #, *p*<0.05; ###, *p*< 0.001.

Since the racemic mixture, N-acetyl-D-L-leucine (Tanganil) has also been suggested to have beneficial effects in lysosomal storage diseases (Kaya et al., 2020), we examined the effects of the separate enantiomers levacetylleucine and N-acetyl-D-leucine as well as the racemate N-acetyl-D-L-leucine on TFEB translocation to the nucleus (**Fig. 3**). While levacetylleucine caused a robust translocation (activation) of TFEB into the nucleus, the D-enantiomer, N-acetyl-D-leucine, had no effect. Remarkably, the racemate N-acetyl-D-L-leucine also failed to activate TFEB translocation, indicating that the presence of the D-enantiomer in the racemic mixture antagonises the effect of the pharmacologically active L-enantiomer, levacetylleucine.

**Fig. 3.**
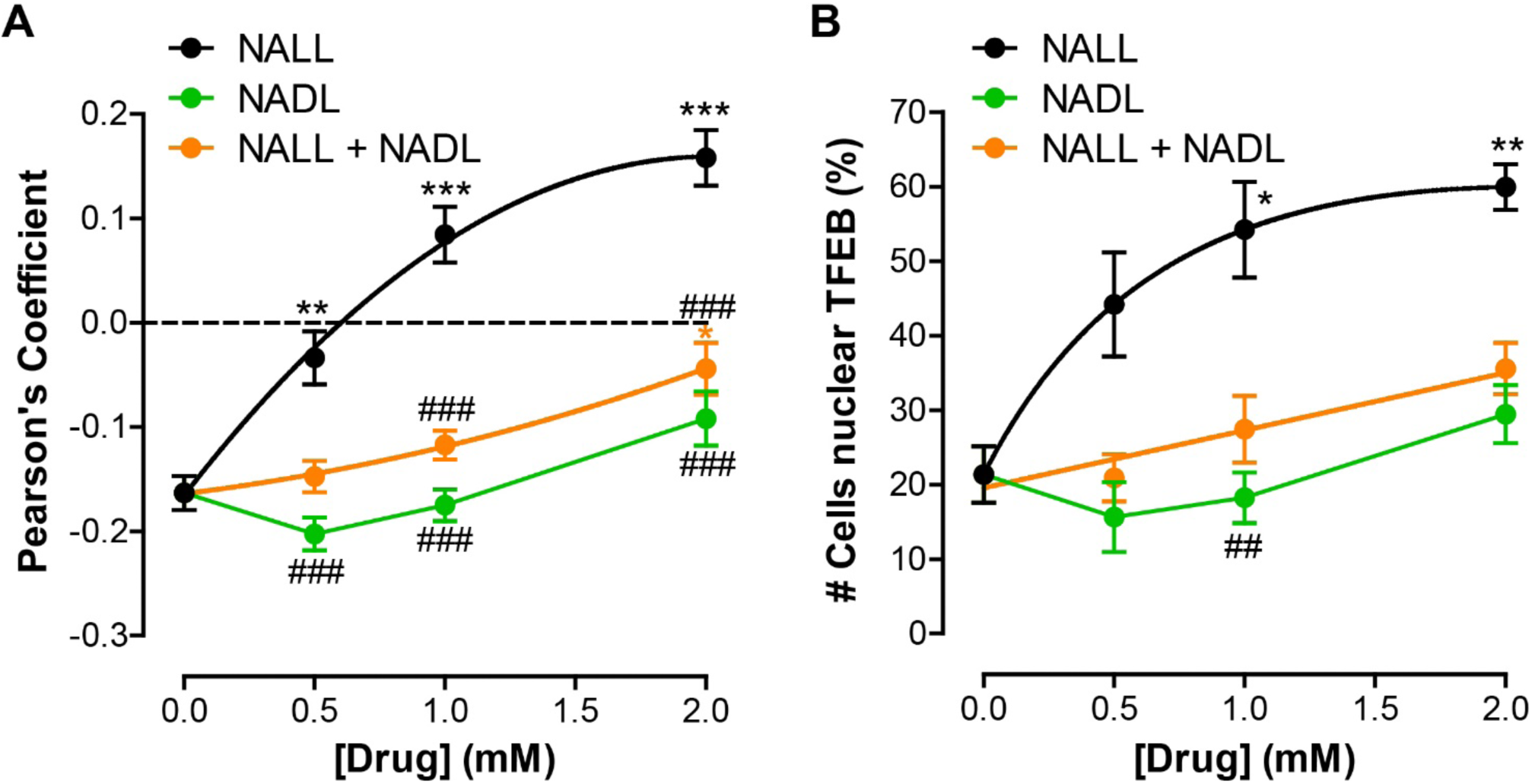
NALL is the most efficacious enantiomer to evoke TFEB translocation. TFEB-EGFP translocation in HeLa cells incubated for 18 hrs with different concentrations of each enantiomer: the L-form (NALL, black), D-form (NADL, green) or the racemic mixture of both (orange). Translocation is expressed either as the Pearson’s correlation coefficient with NucSpot 650 (A) or as the percentage of cells with nuclear TFEB (B). Data are expressed as the mean ± SEM of 153-265 cells, and significance determined by nonparametric ANOVA test, with significance denoted by * *p*< 0.05, ** *p*< 0.01, or *** *p*< 0.001 compared to vehicle control, i.e. 0 mM. Comparing NALL and NADL or racemic mixture, using a nonparametric ANOVA test: ##, *p* < 0.01; ###, *p* < 0.001.

To assess if the activation of TFEB and its nuclear translocation has functional consequences for the HeLa cell, we showed that incubation of cells with levacetylleucine (2 mM) increased the expression of the major lysosomal transmembrane protein, LAMP1 (**Fig. 4**) whose gene has a CLEAR sequence in its promoter region which affords regulation of its expression by TFEB (Palmieri et al., 2011).

**Fig. 4.**
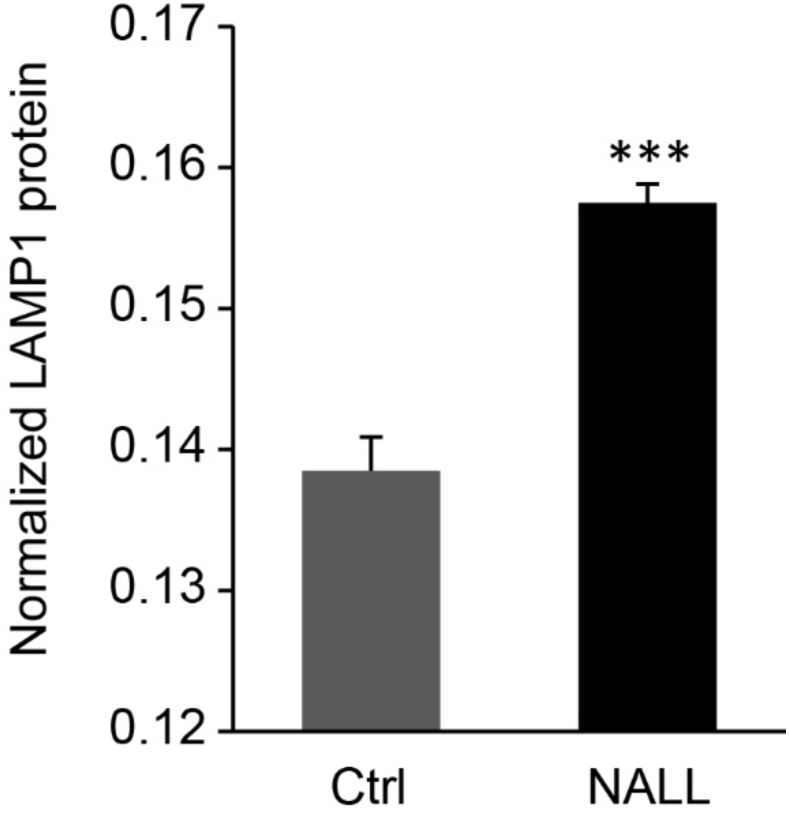
NALL increases LAMP1 protein expression. HeLa cells were treated with 2 mM NALL for 18 hrs, then processed for In-Cell Western analysis: endogenous LAMP1 was labelled with an anti-LAMP1 antibody and stained with CellTag700 (for normalization to cell number). Data are expressed as the mean (LAMP1/CellTag700) ± SEM of 12 replicates, and significance determined by a paired t-test, with significance denoted by *** *p* < 0.001 compared to vehicle control (Ctrl), i.e. 0 mM NALL.

### Effects in NPC1^-/-^ cells

These experiments were then replicated in CRISPR-edited NPC1^-/-^ cells. In contrast to the predominantly cytoplasmic distribution in untreated wild-type HeLa cells, we found that in CRISPR-edited NPC1^-/-^ cells there was significantly higher TFEB localisation (**Fig. 5**), e.g. the overactivation of TFEB activity. We also found that incubation of wild-type cells with the NPC1 small molecule inhibitor, U18666A (Lu et al., 2015) increased the nuclear translocation by a similar degree to 2 mM levacetylleucine (**Fig. 5**).

**Fig. 5.**
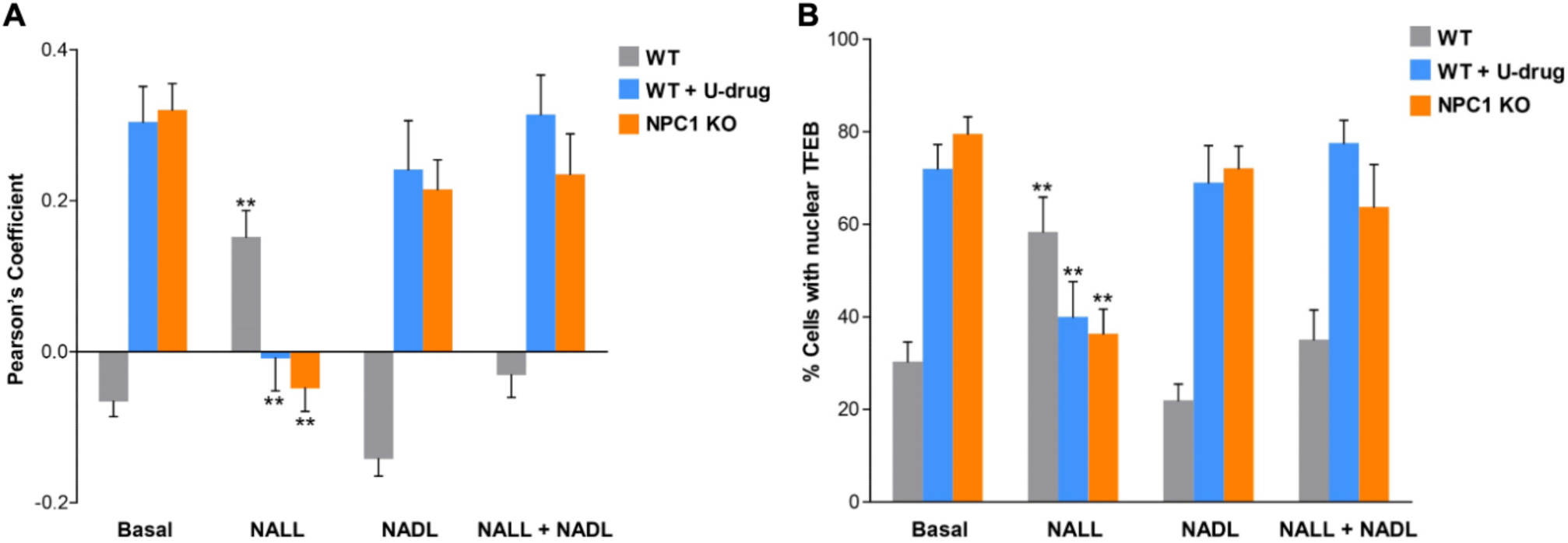
TFEB is constitutively active in NPC and reduced by NALL, but not by NADL or the racemic NADLL. WT HeLa, WT HeLa treated with 2 µg/ml U18666A (U-drug) for 22 hrs and NPC1 KO HeLa cells were transfected with TFEB-mScarlet3 and then incubated with 2 mM L-form (levacetylleucine (NALL), 2 mM D-form (N-acetyl D-leucine (NADL), or the racemic mixture of both (N-acetyl D-L-leucine (NALL + NADL), 1 mM of each) for 22 hrs. Cells were then counter-stained with the NucSpot 488 to label the nucleus and TFEB translocation quantified by colocalization with NucSpot. Nuclear localization is plotted as either the Pearson’s coefficient (A) or derived as the percentage of cells with nuclear TFEB (B). Data are expressed as the mean ± SEM of 101-321 cells, and significance determined by ANOVA test, with significance depicted as ** *p* < 0.01, compared to respective basal control.

In NPC1^-/-^ cells and U18666A-incubated wild-type cells, treatment with levacetylleucine remarkably corrected this situation by reducing nuclear localization to a level around that seen with treatment of wild-type cells with levacetylleucine (2 mM). This down-regulation of TFEB in cells where overstimulated reflected a normalization of TFEB localisation, and was also stereo-specific as for TFEB activation in wildtype cells (Fig. 2), in that the D-enantiomer or racemate had no effect (**Fig. 5**).

## Discussion

The major findings from this study were that levacetylleucine has a biphasic action on TFEB translocation. Given that L-leucine is a key activator of mTOR, the major kinase phosphorylating TFEB (Settembre et al., 2012), it would be expected that levacetylleucine decreases TFEB translocation to the nucleus. In contrast, we have shown a biphasic activity that works to achieve cellular homeostasis depending on the dysfunctional pathophysiology driving the disease: levacetylleucine can stimulate TFEB translocation to the nucleus when it is deficient, enhancing lysosomal biogenesis and autophagy. This effect could serve to ameliorate lysosomal storage disorders and promote autophagy and potentially lysosomal exocytosis, and may be a potential facet of levacetylleucine’s mechanism of action in NPC. However, uniquely, levacetylleucine also downregulates TFEB in cells where it is overstimulated, thereby maintaining cellular homeostasis. This biphasic action is essential for a therapeutic effect **and** to ensure TFEB is not persistently hyper-active (**Fig. 5**). While TFEB activation is often linked to the clearance of protein aggregates and damaged organelles, its sustained and excessive activity can disrupt cellular processes, promote tumour growth, and even induce neuronal cell death. Indeed, the small molecule inhibitor of TFEB, Eltrombopag (Lin et al., 2023) inhibits excessive autophagy by binding to the basic helix-loop-helix-leucine zipper domain of TFEB and has been proposed for repurposing use as a tumour therapeutic. Levacetylleucine likely only transiently activates TFEB due to its rapid metabolism to L-leucine (Churchill et al., 2020), mitigating against the effects of excessive or chronic TFEB stimulation.

These data demonstrate that levacetylleucine acts as a bidirectional modulator of TFEB with normalization of TFEB activation in NPC1^-/-^ cells to levels that may be optimal for correct lysosomal function, whilst in cells with low TFEB activity, levacetylleucine increases it. It could even be thought of as a partial agonist (Ohlsen and Pilowsky, 2005) in the pathways that lead to TFEB activation, with the ability to enhance TFEB activation, whilst antagonising excessive activation as seen in some disease states. This may be a major mechanism that explains levacetylleucine’s neuroprotective effects in NPC and for other neurological, neurodegenerative and neurodevelopmental diseases, and is also directly consistent with levacetylleucine’s biphasic action on mitochondria function (see below) and normalisation of neuronal membrane potential and excitability (Vibert and Vidal, 2001).

We have previously shown that levacetylleucine has a biphasic effect normalising mitochondrial function; levacetylleucine rebalances energy metabolism, irrespective of whether the pathophysiology of the disease is characterized by insufficient or excessive Krebs cycle flux, changing the expression levels of key enzymes that regulate these key pathways (Kaya et al., 2020; Kaya et al., 2021).

The efficacy of the drug is dependent on the acetyl moiety, which renders the drug a substrate for high-capacity MCT transporters and is delivered to cells in supraphysiological quantities and therein transformed into L-leucine by acylases, which we have shown to be expressed in HeLa cells. Such high concentrations of L-leucine would have been expected to stimulate mTOR (Wolfson et al., 2016), and in consequence phosphorylate TFEB and reduce its transcriptional activity as we see in NPC1^-/-^ cells (Fig. 5). In contrast, another acetylated analogue of leucine, N-acetyl-leucine amide has been reported to inhibit mTOR (Hidayat et al., 2003) and this effect may be shared by levacylleucine prior to its metabolism to L-leucine and this could potentially lead to activation of TFEB as we see in wildtype cells (Fig. 1). A transient inhibition may be sufficient to suppress mTOR-mediated TFEB phosphorylation to allow sufficient TFEB translocation to the nucleus to activate the CLEAR network of lysosomal and autophagy genes as we have demonstrated for LAMP1 expression (**Fig. 4**) as an exemplar.

The effects of levacetylleucine that we have observed here occur at concentrations that approximate plasma concentrations during therapeutic dosing (Churchill et al., 2020). Specifically, our findings demonstrate that levacetylleucine (Aqneursa) at therapeutically relevant concentrations significantly normalises TFEB activity improving lysosomal biogenesis and function. Of note, TFEB activation has recently been proposed as a mechanism of action for another newly approved NPC drug, arimoclomol (Shammas et al., 2025). However, (1) despite the findings that TFEB is constitutively active in NPC, arimoclomol had the limited ability to only activate TFEB (as opposed to normalise or down-regulate TFEB), potentially problematic as TFEB’s persistent overexpression can excessively and detrimentally activate autophagy and lysosomes, leading to disruptions in essential cellular processes due to energy strains, and irreversible autophagy-induced neuronal damage and (2) the nominal effects on TFEB activation with arimcolomol in NPC cells were only seen at several orders of magnitude of the concentrations associated with therapeutic dosing (Cudkowicz et al., 2008).

Another important finding was that the effect of levacetylleucine was stereospecific in that N-acetyl-D-leucine did not activate TFEB nuclear translocation (**Fig. 3**). Remarkably, the racemate, N-acetyl-D,L-leucine, was also without effect, indicating that the presence of equimolar N-acetyl D-leucine inhibited the pharmacological activity of the L-enantiomer. Since N-acetyl D,L-leucine, which is sold as the vertigo treatment, Tanganil (Vibert and Vidal, 2001), lacks effect on an important lysosomal signalling pathway which improves lysosomal function, this would correlate with a significantly greater efficacy of levacetylleucine over Tanganil for the treatment of lysosomal storage diseases.

## Conclusions

The goal of this study was to increase our understanding of the cellular mechanisms of action of levacetylleucine. This drug, marketed under the tradename of Aqneursa, underwent a successful phase III trial for NPC patients (Bremova-Ertl et al., 2024), and was approved in September 2024 by the FDA as a stand-alone therapy for NPC (Mullard, 2024). The drug significantly improves both gait and cognitive impairments in these patients within a few weeks, and this clinical improvement is maintained following 18 months of therapy, demonstrating disease-modifying, neuroprotective effects (Patterson et al., 2025).

Here, we have shown that levacetylleucine normalizes the translocation of TFEB to the nucleus, and importantly that this effect was not seen with the amino acid L-leucine, or with the D-enantiomer of N-acetyl leucine. Levacetylleucine’s unique, biphasic ability to stimulate or down-regulate TFEB nuclear translocation and lysosomal function indicates its potential utility in treating a broad range of neurodegenerative and neurodevelopmental disorders characterized by lysosomal and mitochondrial dysfunction. For example, increases in TFEB activity have been suggested to enhance the degradation of Aβ plaques in mouse hippocampus by 40% (Xiao et al., 2015), which may be relevant for human disease since in preclinical models, a 25% Aβ reduction correlates with a meaningful improvement in cognitive deficits (Singh et al., 2024). It has also been shown that TFEB activation promotes the degradation of Tau proteins, leading to a reduction in tau aggregates (Polito et al., 2014) and that a small molecule activator of TFEB ameliorates beta-amyloid precursor protein and tau pathology in Alzheimer’s disease models (Song et al., 2016). Studies have confirmed that TFEB can reduce the deposition of toxic tau aggregates, which represents a neuroprotective effect. TFEB enhances the lysosomal degradation of tau, thereby reducing the formation of toxic aggregates. These mechanisms are crucial for the prevention of tauopathies, such as those observed in Alzheimer’s and other neurodegenerative diseases.

TFEB has been shown to have a protective effect against α-synuclein aggregates, which are typically associated with Parkinson’s disease and Multiple System Atrophy (MSA). TFEB promotes autophagy and the lysosomal degradation of α-Synuclein oligomers, which can reduce the toxic effects of these aggregations (Decressac et al., 2013). In animal models, TFEB overexpression resulted in nearly complete prevention of dopaminergic neuronal loss, indicating the neuroprotective effect of TFEB. Moreover, TFEB overexpression in a mouse model increases dopamine release by 20%, reverses the atrophy of dopaminergic neurons, and enhances protein biosynthesis through mTORC1 phosphorylation (Torra et al., 2018). Consistent with this, the effects of levacetylleucine on an alpha-synucleinonpathy, Parkinson’s disease and its prodromal stage, REM sleep behaviour disorder (RBD) was recently demonstrated in a case series: N-acetyl leucine improved symptoms, reversed loss of striatal dopamine-transporter binding and stabilized a pathological metabolic brain pattern (Oertel et al., 2024); a recent study has also shown that levacetylleucine lowers pS129-synuclein and improves synaptic function in models of Parkinson’s disease (Song et al., 2025).

Our new findings show that levacetylleucine, an approved treatment for NPC, activates the TFEB pathway (scheme: **Fig. 6**), and has a biphasic action: by entering key metabolic pathways and subject to complex feedback mechanisms in signalling pathways, it normalizes TFEB activity in disease cells thereby restoring normal cellular homeostasis, (Kaya et al., 2021) reinforcing its tremendous promise as a new therapy for broad neurodegenerative and neurodevelopmental disease management.

**Fig. 6.**
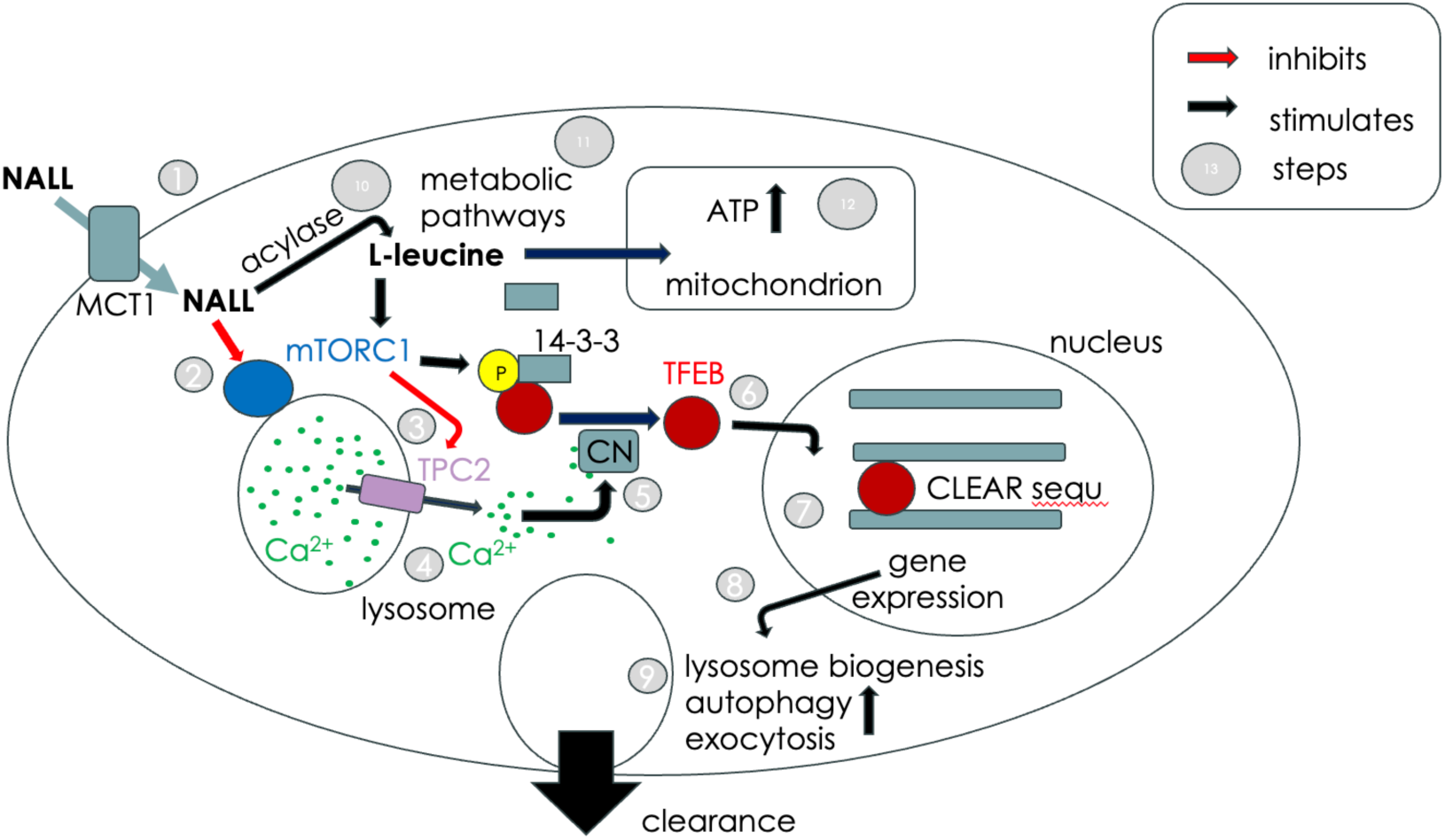
Scheme for the mechanisms of action of levacetylleucine (NALL) to enhance lysosomal and mitochondrial function. **Step 1** Levacetylleucine (NALL) is transported into cells via the MCT transporter family. **Step 2** Levacetylleucine inhibits the activity of the mTORC complex and reduces TFEB phosphorylation. **Step 3** Inhibition of mTOR relives its inhibition on Ca^2+^ release channels expressed in lysosomes leading to Ca^2+^ release by TPC2 and/or TRPML1 (**Step 4**). **Step 5** Lysosomal Ca^2+^ release activates calcineurin (CN) to dephosphorylate TFEB and remove its ability to bind 14-3-3 proteins allowing it to translocate to the nucleus (**Step 6**). **Step 7** TFEB bind to CLEAR gene promoters and activates lysosomal and autophagic gene expression. **Step 8** This increases lysosomal biogenesis, autophagy and lysosomal exocytosis leading to cellular clearance of stored lysosomal material (**Step 9**). **Step 10** Levacetylleucine is metabolised to L-leucine that reactivates mTOR resulting in TFEB inactivation through phosphorylation. **Step 11** L-leucine enters metabolic pathways, where it increases mitochondrial ATP production (**Step 12**).

## Abbreviations

ATP: adenosine triphosphate
CLEAR: network coordinated lysosomal expression and regulation network
ER: endoplasmic reticulum
FDA: Food and Drug Administration (USA)
INN: International Nonproprietary Names
LAT: L-type amino-acid transporter
LSD: lysosomal storage disease
MCT: monocarboxylate transporter
mTORC1: mammalian target of rapamycin complex 1
NPC: Niemann-Pick disease type C
NALL: N-acetyl-L-leucine
MCS: membrane contact site(s)
RBD: REM sleep behaviour disorder
TFEB: Transcription Factor EB
USAN: United States Adopted Name

---

**Fig. S1.**
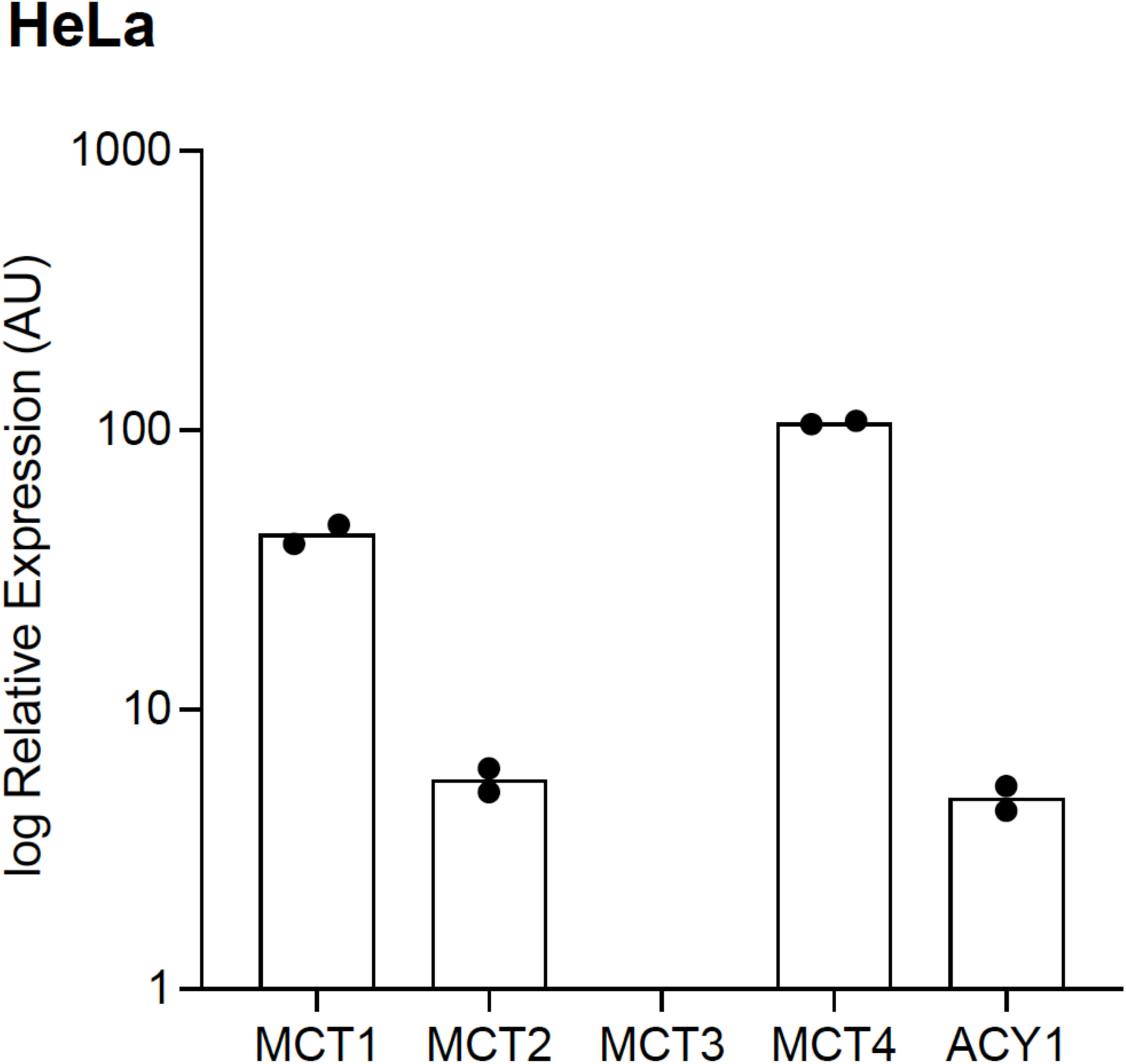
Expression patterns of monocarboxylate transporters (MCT1, MCT2, MCT3, MCT4) and the aminoacylase 1 enzyme (ACY1). These were determined by RT-qPCR. Expression was normalised to β -actin, on a scale where β-actin expression equals 10,000 arbitrary units. Data are shown as the mean (bars) of two biological replicates (symbols). MCT3 expression was not detected.

